# Patchy distribution of a Madrean Sky Island squirrel shaped by historical habitat configuration

**DOI:** 10.64898/2026.02.23.707456

**Authors:** Binaya Adhikari, Jesse M Alston, Joseph R Burger

## Abstract

Sky islands, mountain-top forests isolated by surrounding lowlands, offer unique opportunities to test how past and present landscapes shape species distributions. We examined the distribution of the Arizona gray squirrel (*Sciurus arizonensis*) across the Madrean Archipelago to test the constraint-based dynamic island biogeography (C-DIB) model, which posits that current occupancy in the sky islands reflects historical habitat size and connectivity. Using verified specimen records, we modeled climatically suitable habitats across four time periods: the Last Glacial Maximum (LGM), Mid-Holocene (MH), Present, and Future. For each mountain, we quantified suitable habitat area and estimated least-cost dispersal distances to assess both persistence and colonization potential. Our results suggest that species presence is best explained by LGM habitat metrics, which marginally outperformed models based on current conditions. Mountains that were large or well-connected during the LGM continue to support *S. arizonensis*, whereas historically isolated ranges remain unoccupied despite suitable contemporary habitat. These findings indicate a legacy of Pleistocene connectivity and reveal patterns of distributional disequilibrium. Furthermore, climatically suitable habitat for *S. arizonensis* has shifted both elevationally and geographically through time, reflecting long-term responses to climatic change. Together, these results emphasize the importance of protecting historically connected refugia, restoring riparian corridors that facilitate dispersal, and developing mountain range-specific management strategies that account for elevational shifts and potential downslope habitat recovery under future climate scenarios.

---

Sky islands are montane ecosystems rising above surrounding lowlands with distinct climates, habitats, and biota. In the Madrean region of southeastern Arizona, southwestern New Mexico, and northern Sonora and Chihuahua, these isolated ranges form a mosaic of forested uplands surrounded by desert basins. Their steep environmental gradients and isolation create unique ecological “islands” that harbor relict and endemic species (DeBano et al. 1995; López-Hoffman and Quijada-Mascareñas 2012; Yanahan and Moore 2019). Because they differ sharply from the surrounding landscape yet remain close enough for occasional dispersal, sky islands provide an exceptional setting for studying how habitat configuration and historical connectivity shape present-day species distributions (Wiens et al. 2019; Cheng et al. 2024).

Climatic oscillations throughout the Quaternary have shaped biogeographic patterns in these systems (Davies et al. 2011; Favé et al. 2015; Wiens et al. 2019; Love et al. 2023). Today, rapid anthropogenic climate change is similarly transforming biodiversity worldwide by altering temperature and precipitation regimes, shifting species ranges, and fragmenting suitable environments, with especially pronounced effects in montane ecosystems (Bhandari et al. 2022; Adhikari et al. 2023; Baral et al. 2023, 2024; Subedi et al. 2023). Island area has long been recognized as a key determinant of persistence in insular populations (Brown 1971; Lomolino et al. 1989; MacArthur and Wilson 2001). Larger islands can support greater habitat diversity and larger populations, thereby reducing extinction risk even when connectivity is limited (Lomolino et al. 1989). In the Madrean system, the extent of forested habitat during glacial periods may have been more critical for long-term persistence than dispersal itself, which could have primarily served to rescue small populations. During the cooler, wetter Pleistocene, oak and pine woodlands extended into lower elevations, creating connectivity among many mountain ranges(McCormack et al. 2008). This enhanced connectivity, especially during the Last Glacial Maximum (LGM, ∼21 ka), facilitated movement of forest-dependent species (Ramírez-Barahona and Eguiarte 2013; Nekola et al. 2024). In contrast, the warmer, drier Mid-Holocene (MH, ∼6 ka) drove these corridors upslope, fragmenting woodlands and isolating populations (Cain et al. 1998; Wiens et al. 2019). Consequently, contemporary distributions in the Madrean Sky Islands reflect intertwined legacies of historical area and connectivity, which together continue to structure regional biodiversity patterns.

The Equilibrium Theory of Island Biogeography (MacArthur and Wilson 1967) proposed that species richness on islands reflects a dynamic balance between colonization and extinction, shaped by island area and isolation. Larger and better-connected islands tend to support more species, a pattern long extended to terrestrial habitat islands such as the Madrean ranges (Brown 1971; Lomolino et al. 1989; Herrera et al. 2017). The Constraint-Based Dynamic Island Biogeography (C-DIB) model (Burger et al. 2019) builds upon this foundation by integrating both historical and contemporary landscape structure. It highlights legacy effects or hysteresis, where populations established during past periods of connectivity may persist despite current isolation or marginal habitat conditions. This perspective emphasizes the need to consider temporal context when interpreting present-day patterns of persistence in sky island systems.

Tree squirrels (*Sciuridae*) offer an excellent model for studying sky island biogeography because their distributions integrate both habitat constraints and historical dispersal. The Arizona gray squirrel (*Sciurus arizonensis*), a riparian specialist, exhibits a fragmented distribution across the Madrean Sky Islands, largely limited by the extent of riverine corridors through surrounding deserts (Pasch and Koprowski 2005). In some ranges, squirrels are absent despite apparently suitable habitat, suggesting that historical barriers restricted colonization; in others, small patch size and human pressures have likely driven local extinctions (Koprowski 2005; Pasch and Koprowski 2011; Abreu-Jr et al. 2020). Comparable patterns occur in related taxa: a population of the Mexican fox squirrel (*S. nayaritensis*) persists only in the Chiricahua Mountains north of Mexico, the Mount Graham red squirrel (*Tamiasciurus fremonti*) persists solely in the Piñaleno Mountains, and Abert’s squirrel (*S. aberti*), native to the Rockies and Sierra Madre Occidental, occurs in the Madrean region only through twentieth-century introductions (Hutton et al. 2003; Edelman and Koprowski 2005). Collectively, these contrasting distributions underscore how historical habitat area and shifting connectivity have shaped present-day patterns of persistence, with many populations remaining as relicts of past forest expansions but showing little evidence of recent immigration (Brown 1971; Lomolino et al. 1989; Pasch and Koprowski 2005).

Two main factors are expected to influence squirrel presence on a sky island: (1) the area and quality of available habitat (e.g., riparian forests), which determine persistence and extinction risk, and (2) the connectivity of each range, both historical and current, to other occupied mountains, which shapes colonization potential. Larger patches are likely to sustain more stable and genetically diverse populations, whereas isolated ranges may remain unoccupied despite suitable climatic conditions (Scheffer et al. 2006; Prugh et al. 2008; Spiesman, Stapper and Inouye 2018; Huxley and Spasojevic 2021).

We examined how past and present landscape structure the distribution of *S. arizonensis* across the fragmented Madrean Sky Islands within the C-DIB framework. Specifically, we tested two complementary non-mutually exclusive hypotheses: (H1) Temporal legacy: islands with greater climatic-suitability area or higher connectivity during the LGM or Mid-Holocene are more likely to be occupied today, independent of present conditions; and (H2) Structural drivers: current occurrence reflects the additive effects of habitat area (expected β > 0, where β is the regression coefficient describing the expected direction of the effect) and dispersal cost (expected β < 0) integrated across time. Here, “structural” refers to the geometry and separation of habitat patches, measured as climatic suitability–derived area and least-cost connectivity across the LGM, MH, and Present. Using verified specimen records, we contrasted time-partitioned models (H1) with composite, time-integrated models (H2) to test whether modern distributions primarily reflect legacies of glacial refugia or the cumulative influence of area–isolation structure in the archipelago.

## Materials and methods

The Madrean Sky Islands include about 55 mountain ranges spanning southeastern Arizona, southwestern New Mexico (USA), and northern Sonora and Chihuahua (Mexico). For this study, we analyzed 29 U.S. ranges rising above ∼1,500 m that support montane woodland or mixed conifer vegetation (Fig. 1). Within these systems, *S. arizonensis* primarily occupies riparian forests along drainage networks but also uses adjacent pine–oak and mixed conifer woodlands for nesting and foraging (Pasch and Koprowski 2005; Cudworth and Koprowski 2011). We focused on the U.S. ranges, which have been surveyed more extensively, providing greater confidence in presence–absence data.

**Figure 1.**
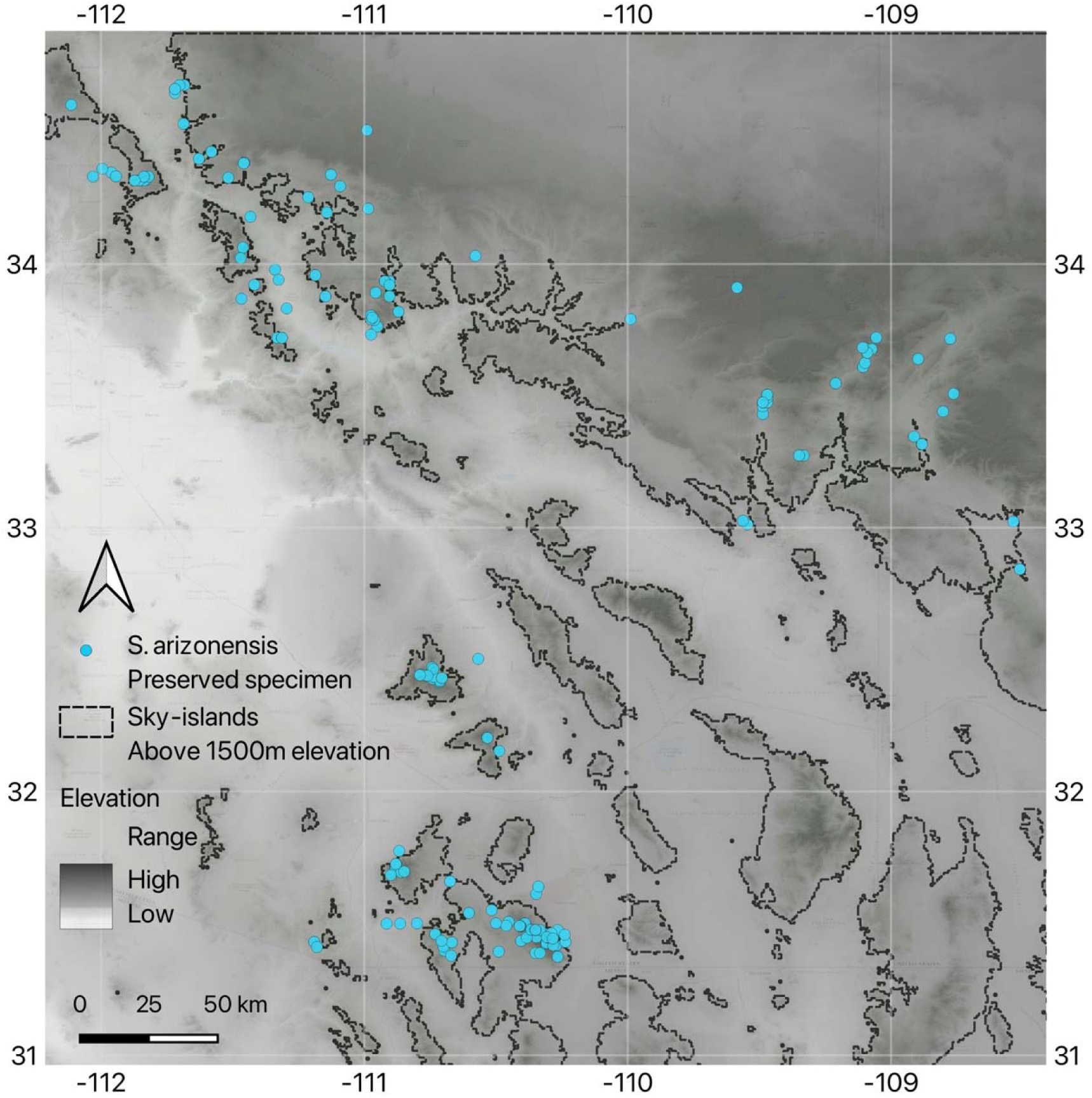
Map showing occurrence records of the Arizona gray squirrel (*Sciurus arizonensis*) across the Madrean Sky Islands. Each blue point represents preserved specimen occurrence from Global Biodiversity Information Facility (GBIF) (https://doi.org/10.15468/dl.5n2m9t). Sky islands are represented as isolated montane areas above 1500 m elevation (black dashed outlines). Elevation is shown in grayscale, with darker tones indicating higher elevations. The map highlights the patchy distribution of *S. arizonensis* across isolated mountain ranges separated by desert lowlands.

We initially compiled occurrence records for native tree squirrels known from the Madrean Sky Islands using verified museum specimen localities with reliable geographic coordinates obtained primarily from the Global Biodiversity Information Facility (GBIF) (Supplementary data: SD1). Each mountain range was classified as either occupied (1) or unoccupied (0) based on verified records. Given the conspicuous behavior of diurnal tree squirrels (vocalizations, nests, feeding signs), the extensive history of biodiversity surveys, we consider established populations to be reliably detected.

Among the regional species, only *S. arizonensis* occurs across multiple U.S. sky islands with sufficient distributional breadth for hypothesis testing. Other species were excluded because their ranges are restricted to single mountains (*S. nayaritensis, T. fremonti*) or reflect recent human introductions (*S. aberti*) (Best 1995; Hutton et al. 2003; Edelman and Koprowski 2005; Koprowski 2005). Focusing on *S. arizonensis* therefore allows inference from naturally distributed populations with adequate spatial coverage, enabling robust assessment of how habitat area and historical connectivity influence species persistence. As a deciduous riparian forest specialists with limited ability to disperse across desert lowlands, *S. arizonensis* represents an ideal model for testing dispersal limitation in montane mammals (Frey et al. 2008; Cudworth and Koprowski 2011).

### Species Distribution Modeling

We modeled the climatic niche of *S. arizonensis* using the Maxnet algorithm implemented in Wallace (Phillips et al. 2006; Kass et al. 2018), with the 19 CHELSA bioclimatic variables at ∼1 km resolution (Karger et al. 2017). From 187 georeferenced museum specimens, we applied a 1-km spatial thinning filter in the *spThin* package (Aiello Lammens et al. 2015) to reduce spatial autocorrelation and sampling bias, retaining 157 unique records (Fig. 1). To focus model calibration on the species’ core climatic niche, we further restricted the dataset to occurrences from the contiguous Rocky Mountain–Colorado Plateau forests (n = 88), excluding records from the Madrean Sky Islands. This approach ensured that the model captured the environmental conditions of the species’ primary range while avoiding bias introduced by peripheral, potentially ecologically divergent populations (Barve et al. 2011; Owens et al. 2013).

All 19 CHELSA bioclimatic variables were initially considered during model tuning. To minimize multicollinearity and retain ecological interpretability, we evaluated Variance Inflation Factors (VIF) and retained five predictors (BIO01, BIO07, BIO12, BIO15, BIO17) with VIF < 6. These variables represent the major temperature and precipitation gradients constraining the species’ riparian–woodland niche: annual mean temperature (BIO01), temperature annual range (BIO07), annual precipitation (BIO12), precipitation seasonality (BIO15), and precipitation of the driest quarter (BIO17) (Supplementary data: SD2). This selection followed established recommendations for ecological niche modeling to balance information retention and parsimony (Zuur et al. 2010; Braunisch et al. 2013; Dormann et al. 2013).

The 88 occurrence records were partitioned randomly into training (70%) and testing (30%) datasets (Joseph 2022). We performed a full hyperparameter search across four feature-class combinations (Linear, Quadratic, Product, and Hinge) and eight regularization multipliers (0.5–3.5, in 0.5 increments) to balance model complexity and predictive performance (Merow et al. 2013; Muscarella et al. 2014). Candidate models were ranked by corrected Akaike Information Criterion (AICc), with ties resolved by maximizing test AUC while maintaining omission rates ≤ 5%, following recommended performance thresholds (Warren and Seifert 2011; Shcheglovitova and Anderson 2013).

The selected model was projected across four climatic periods: the Present (1970–2000), the MH (∼6 ka), the LGM (∼21 ka), and a late-century Future scenario (CMIP6 SSP2-4.5 composite for 2071–2100). These projections represent climatic suitability rather than direct habitat distributions. Because *S. arizonensis* depends on riparian forests that follow stream networks across multiple vegetation zones, our models delineate the broader climatic envelope within which such habitats may occur, without explicitly capturing fine-scale vegetation or hydrological structure. Consequently, results should be interpreted as reflecting climatic constraints on potential habitat, rather than as maps of realized distribution.

### Spatial Analysis of Habitat Patch Characteristics

Spatial analysis of habitat patch characteristics was conducted in R using *terra* (Hijmans et al. 2025) and *landscapemetrics* (Hesselbarth et al. 2019). A climatic suitability raster was generated for each of the four climatic periods and used to quantify changes in the spatial configuration of suitable habitat through time. For each period, we calculated two suitability-weighted metrics for every sky island: (1) mean elevation, representing vertical shifts in optimal habitat, and (2) geographic centroid, denoting the spatial center of suitable pixels. Together, these metrics describe how both the elevational and horizontal positions of favorable habitat have changed across glacial–interglacial cycles, providing a spatial framework for interpreting dispersal pathways and colonization dynamics of *S. arizonensis*.

### Hypothesis Testing Using Area and Connectivity Metrics

To assess whether habitat extent or dispersal connectivity better predicts *S. arizonensis* presence, we developed a workflow for hypothesis testing (Fig. 2). We delineated the 29 sky island ranges of Arizona and New Mexico by applying a 1,500 m elevation contour to a USGS digital elevation model (Fig. 1). This elevation approximates the ecological transition from desert–grassland into pine–oak woodland and mixed conifer forest, consistent with previous studies of Madrean mammals (Frey et al. 2008; Cudworth and Koprowski 2011; Coronel-Arellano et al. 2018). Although *S. arizonensis* occasionally descends into lower-elevation riparian canyons, most verified records occur above this threshold (Frey et al. 2008). For consistency, island boundaries were delineated once using the ≥1,500 m contour and kept fixed across all climatic periods. Period-specific climatic suitability and connectivity metrics were then calculated within these polygons. Each delineated range was treated as a discrete “island,” scored for squirrel presence (1) or absence (0), and characterized by two predictor suites: (1) habitat area and (2) dispersal connectivity.

**Figure 2.**
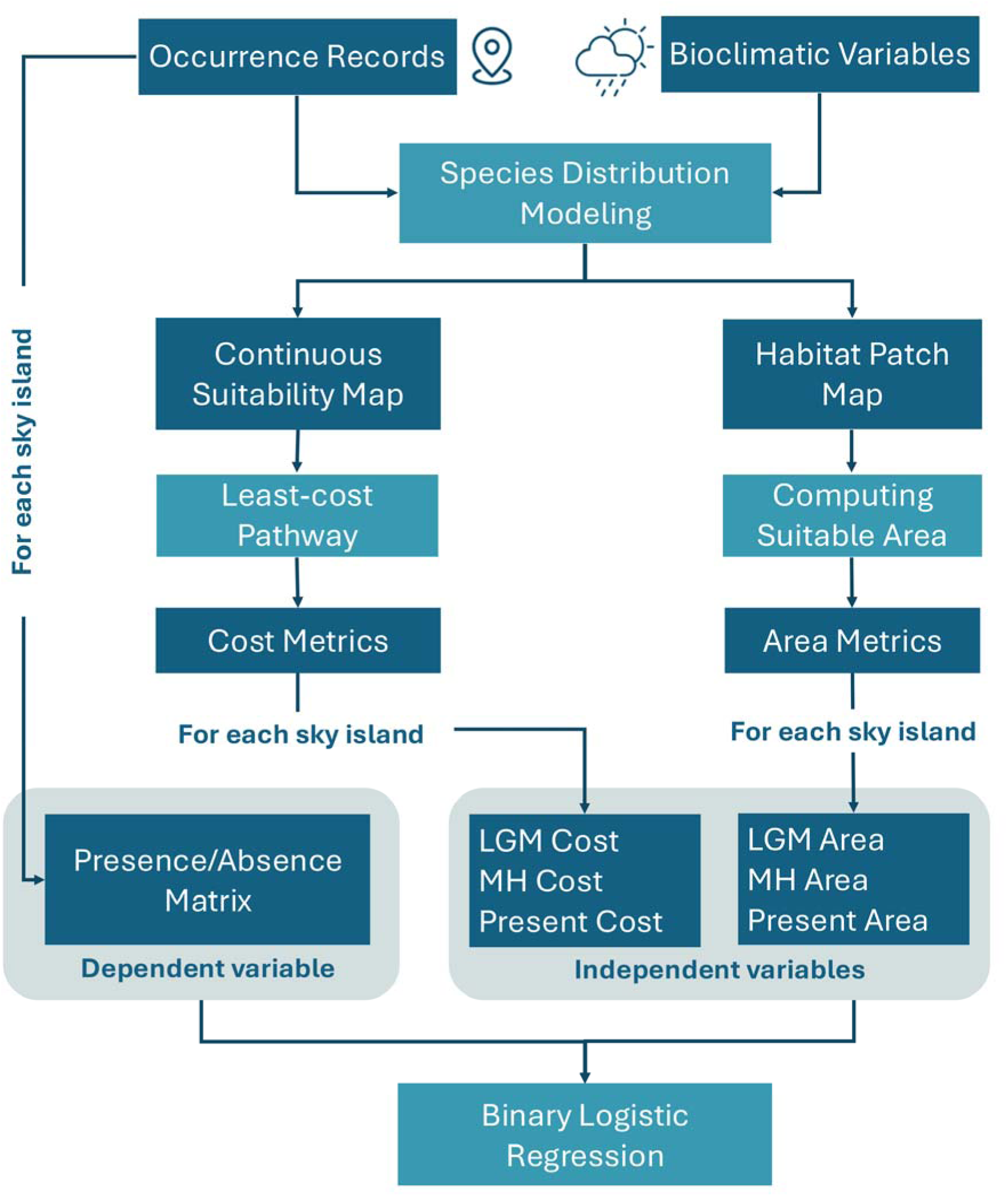
Workflow illustrating the generation of area- and connectivity-based predictors and presence–absence data used to model *Sciurus arizonensis* occupancy across sky islands. Species distribution models developed in Wallace provided habitat suitability maps for the LGM, Mid-Holocene, and Present, from which suitable area and least-cost dispersal cost were derived as predictors in binary logistic regression analyses.

From Maxnet-derived climatic suitability maps for the Present, MH, and LGM, we applied the 10th-percentile training presence threshold (p10) to generate binary suitability layers. Each layer was then clipped to the extent of individual sky islands to calculate period-specific climatically suitable area (km²; Supplementary Data: SD3). The p10 threshold was selected because it provides a balanced trade-off between sensitivity and specificity, minimizes marginal overprediction, and ensures comparability of suitability estimates across time periods (Liu et al. 2005; Jiménez-Valverde and Lobo 2007; Pearson et al. 2007; Liu, Newell and White 2016).

For the connectivity analysis, continuous climatic suitability grids (Fig. 3) were converted to resistance surfaces using the transformation resistance = (1− suitability), where suitability values ranged from 0 to 1. This approach assumes that climatically favorable landscapes impose lower dispersal costs (Adriaensen et al. 2003). Accumulated cost surfaces were computed in QGIS using the GRASS *r.cost* algorithm (GRASS Development Team 2024), quantifying cumulative resistance from mainland source forests on the Mogollon Rim and the Rocky Mountain–Colorado Plateau across the intervening landscape (Etherington and Holland 2013). Using the associated backlink rasters, we then applied the *r.drain* function to delineate the minimum-cost paths (LCPs) between the mainland and the highest-suitability pixel on each sky island (Fig. 4), following standard least-cost connectivity approaches (Adriaensen et al. 2003; Etherington and Holland 2013).

**Figure 3.**
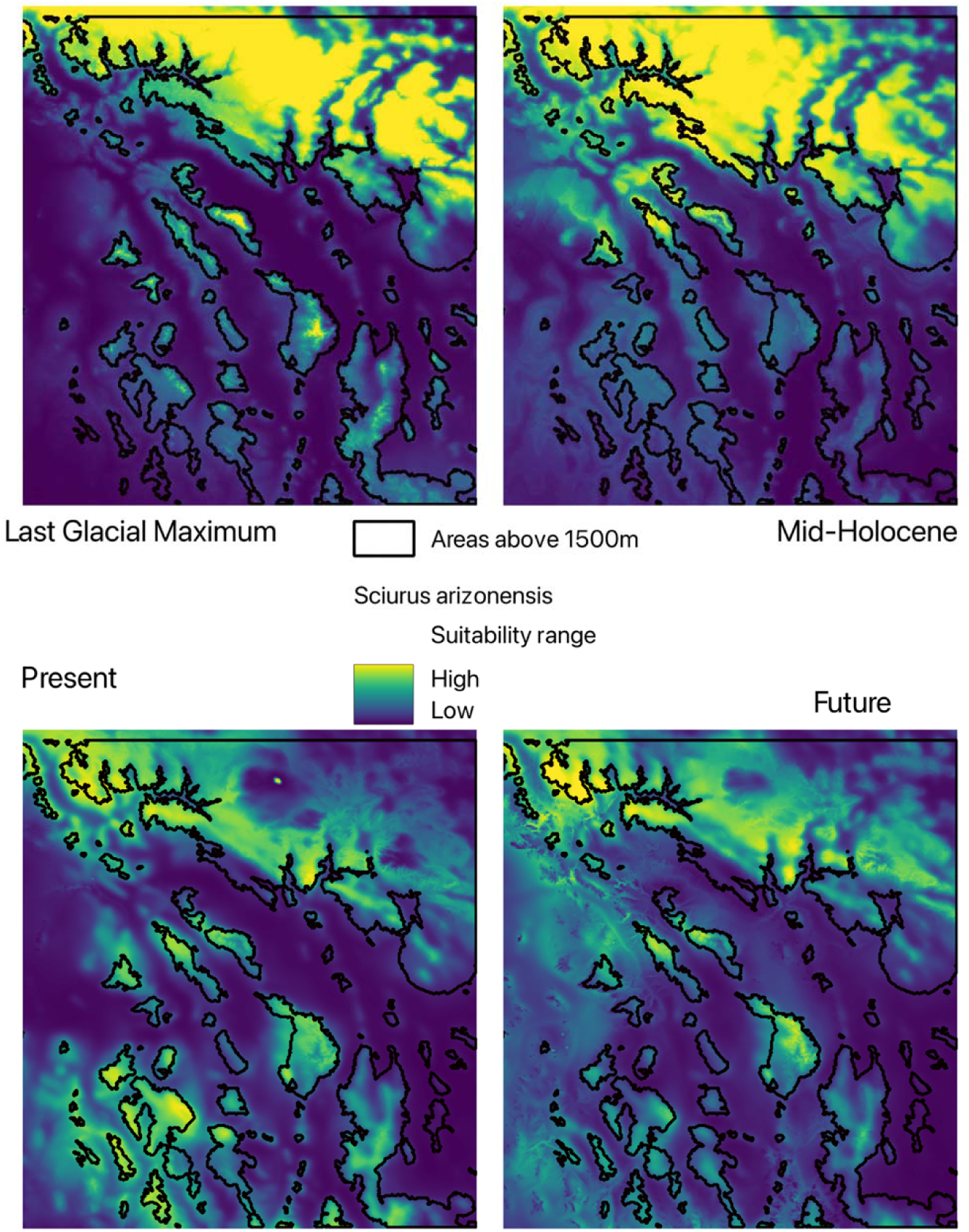
Continuous habitat-suitability maps for *Sciurus arizonensis* across four time slices: Last Glacial Maximum, Mid-Holocene, Present, and a 2070 climate projection. Warmer colours indicate higher suitability, and black outlines mark terrain above 1 500 m. Suitable habitat appears most extensive during the LGM, diminishes and becomes more fragmented by the Mid-Holocene, remains patchy today, and is projected to contract further, with higher-elevation areas retaining the greatest suitability. These patterns suggest a gradual shift from historically connected landscapes toward increasing isolation of sky-island populations.

**Figure 4.**
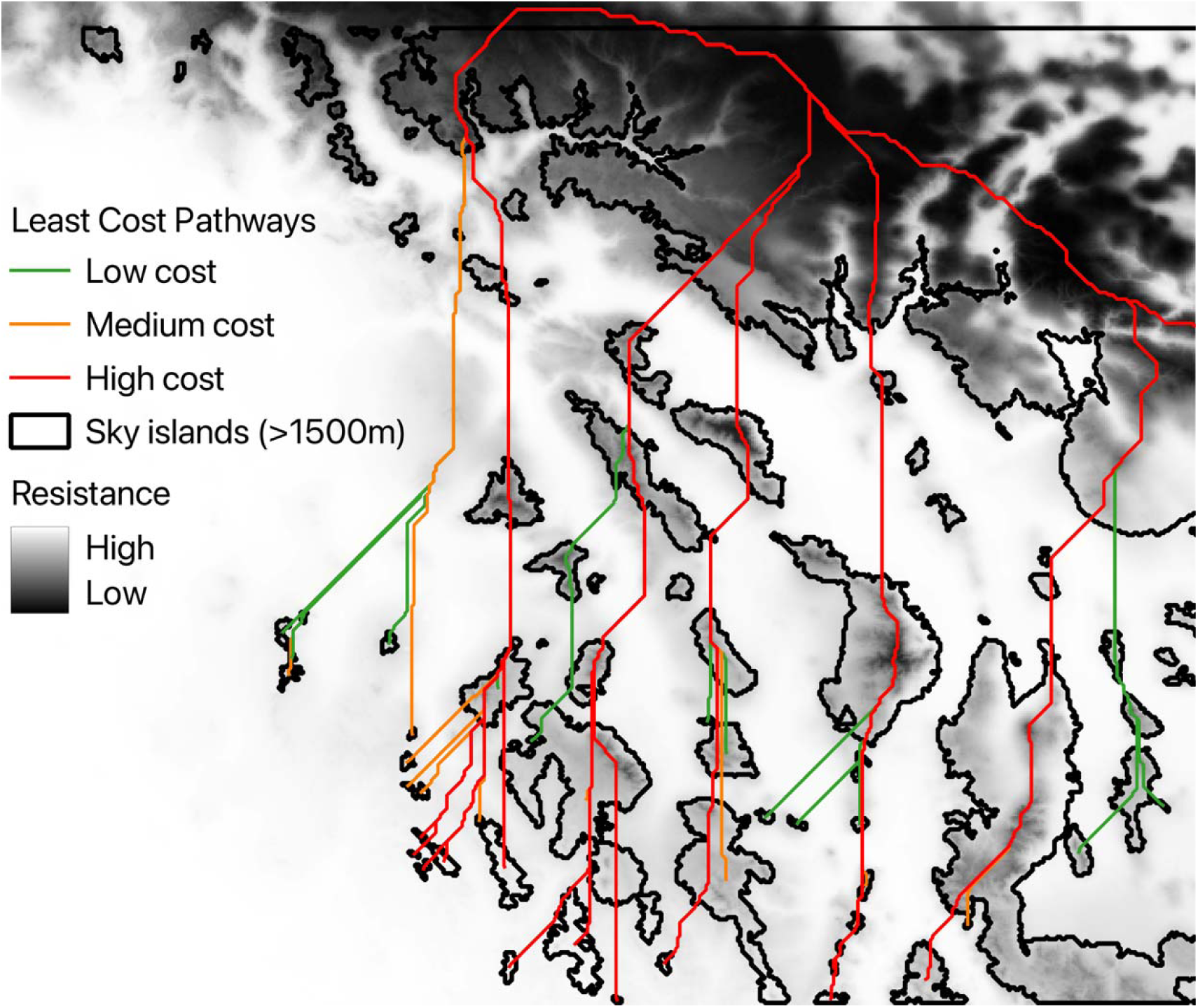
Least-cost pathways connecting the mainland to individual sky islands under Last Glacial Maximum (LGM) conditions, overlaid on a resistance surface derived from the Maxnet climatic-suitability model (resistance = 1 – suitability, range 0–1). Accumulated costs were computed in QGIS using the GRASS *r.cost* and *r.drain* algorithms (8-neighbor). Paths originate from mainland source to the highest-suitability pixel on each island. For visualization purposes, green, orange, and red lines indicate low, moderate, and high cumulative dispersal cost, respectively. Black outlines show fixed island boundaries above 1,500 m.

The resulting LCP values represent the minimum cumulative dispersal costs, where lower values indicate greater historical connectivity and fewer landscape barriers. From these analyses, we derived six predictors: habitat area and LCP distance for each of the three climatic periods (LGM, MH, and Present). These predictors were incorporated into temporal and structural logistic regression models to test the relative importance of habitat extent versus connectivity in shaping *S. arizonensis* distributions across time (Fig. 2).

### Model Building and Statistical Analysis

Given the small sample size (5 presences across 29 sky islands), we modeled *S. arizonensis* occupancy as a binary outcome (presence = 1, absence = 0) using Firth’s penalized logistic regression, implemented in the *logistf* package in R (Heinze et al. 2025). Here, occupancy refers to verified species presence on a sky island based on curated specimen records, representing long-term population persistence rather than detection-corrected probabilities estimated in formal occupancy models (MacKenzie et al. 2002). Traditional logistic regression was unsuitable due to the risk of biased estimates and separation errors common in sparse, unbalanced datasets. Firth’s correction reduces small-sample bias in maximum likelihood estimation and provides stable parameter estimates even when separation occurs.

### Predictor Preparation and Transformation

Predictor variables included habitat area (km²) and least-cost path (LCP) distance, a proxy for dispersal connectivity, for three time periods: LGM, MH, and Present. All continuous predictors were log-transformed using log (x + 1) to reduce skewness and standardize scales across models.

Multicollinearity was assessed using VIF, which indicated strong correlations among area variables and among cost variables across temporal periods, with VIF values exceeding conservative thresholds (VIF > 10). To address this, we employed complementary strategies, including penalized logistic regression, time-partitioned temporal models, and Principal Component Analysis (PCA)-based structural models.

1. Temporal-legacy models (H1)

To evaluate whether historical or contemporary conditions better explain *S. arizonensis* occupancy, we fit three period-specific models, each incorporating area and cost from a single time slice. Predictors were entered independently with expected effects of β_area_ > 0 and β_cost_ < 0, acknowledging that support for H1 does not require both to be significant simultaneously.

LGM model: Occupancy ∼ log (LGM area) + log (LGM cost)

MH model: Occupancy ∼ log (MH area) + log (MH cost)

Present model: Occupancy ∼ log (Present area) + log (Present cost)

This design isolates the predictive strength of each period while minimizing cross-period collinearity.

2. Structural, time-integrated models (H2)

To test whether cumulative landscape structure better predicts occupancy, we used PCA to reduce collinearity among temporally correlated predictors. Separate PCAs were conducted for area and cost variables:

Area PCA inputs: log (LGM area), log (MH area), log (Present area)

Cost PCA inputs: log (LGM cost), log (MH cost), log (Present cost)

From each PCA, the first principal component (PC1) was extracted to represent a composite gradient of habitat area (larger across time) and isolation (lower across time). The resulting additive model tested the integrated effects of these orthogonalized gradients:

Occupancy ∼ PC1_area + PC1_cost (expected signs: PC1_area > 0; PC1_cost < 0)

This approach retains the shared temporal signal of correlated predictors while producing independent axes that capture long-term area–isolation structure, allowing direct evaluation of additive or compensatory effects.

### Model Evaluation and Comparison

Model performance was evaluated using multiple complementary metrics. Model parsimony was assessed with Akaike’s Information Criterion corrected for small sample sizes (AICc) (Hurvich and Tsai 1989), calculated using the MuMIn package in R (Bartoń 2025) following the information-theoretic framework of (Anderson and Burnham 2002). Discriminatory ability was quantified using the area under the receiver operating characteristic curve (AUC) ((Hanley and McNeil 1982) and predictive generalizability was assessed via 5-fold cross-validated AUC (CV-AUC) implemented with the createFolds function in the *caret* package (Kuhn 2008). Model calibration and overall accuracy were evaluated using the Brier score, calculated as the mean squared difference between predicted probabilities and observed outcomes (Brier 1950).

For each model, we reported coefficient estimates, odds ratios, and 95% confidence intervals. All logistic regressions employed Firth’s bias-reduction method (Firth 1993; Heinze and Schemper 2002), implemented via the *logistf* package in R (Heinze et al. 2025), to minimize small-sample bias and mitigate separation issues.

## Results

### Climatic-suitability models

The selected Maxnet model (LQHP, β = 1) performed strongly (test AUC = 0.87–0.90; omission < 3%), demonstrating robust discriminatory ability under spatial cross-validation. BIO12 (annual precipitation; ∼31%) and BIO01 (mean annual temperature; ∼24%) were the dominant contributors, followed by BIO07, BIO15, and BIO17. Response curves (Supplementary Data: SD4) showed increasing climatic suitability with warmer mean temperatures and intermediate precipitation (∼600 mm), patterns consistent with the species’ riparian association and drought sensitivity. These projections were subsequently converted into suitability-based area estimates within each sky island and time slice to test the temporal (H1) and structural (H2) hypotheses.

### H1 - Temporal-legacy (time-partitioned models)

Among the temporal models, the LGM model showed the strongest support (AICc = 28.1; ΔAICc = 0.0; AUC = 0.842; Brier = 0.117), followed closely by the Present model (AICc = 28.2; ΔAICc = 0.1; AUC = 0.758) (Table 1). In the LGM model, habitat area was positively associated with present-day occupancy (β = 0.80; OR = 2.23; 95% CI: 0.86–6.58; p = 0.103), whereas dispersal cost was negatively associated (β = −1.13; OR = 0.32; 95% CI: 0.04–1.06; p = 0.063) (Table 2). These relationships suggest that persistence is more likely in ranges that supported extensive, well-connected glacial refugia, while isolation during the LGM reduced persistence probability.

**Table 1.** Comparison of logistic regression model performance across temporal and structural predictors of *S. arizonensis* occupancy. The LGM model (log-transformed area and cost) had the lowest AICc and highest AUC, suggesting better fit and discriminative power than models based on Mid-Holocene, Present, or PCA-derived structural predictors. CV-AUC and Brier scores provide additional measures of predictive reliability.

| Model type | Model formula | AICc | AUC | CV-AUC | Brier |
| --- | --- | --- | --- | --- | --- |
| Temporal – LGM | log(LGM area) + log(LGM cost) | 28.1 | 0.842 | 0.717 | 0.117 |
| Temporal – MH | log(MH area) + log(MH cost) | 33.5 | 0.567 | 0.783 | 0.143 |
| Temporal – Present | log(Present area) + log(Present cost) | 28.2 | 0.758 | 0.733 | 0.120 |
| Structural | PC1_area + PC1_cost | 30.7 | 0.725 | 0.533 | 0.131 |

**Table 2.** Summary of parameter estimates from temporal and structural Firth logistic regression models explaining *S. arizonensis* occupancy. Predictors include log-transformed area and cost at LGM and Present, and principal components (PC1) derived from area and cost across all time periods. Coefficients, odds ratios (OR), 95% confidence intervals (CI), and p-values are reported. While no predictors were statistically significant at α = 0.05, LGM cost showed a marginally significant negative effect, and LGM area a positive trend.

| Model | Predictor | Coefficient | Odds Ratio<br>(OR) | 95% CI for<br>OR | p-value |
| --- | --- | --- | --- | --- | --- |
| Temporal | log(LGM area) | 0.80 | 2.23 | 0.86 – 6.58 | 0.103 |
| Temporal | log(LGM cost) | -1.13 | 0.32 | 0.04 – 1.06 | 0.063 |
| Temporal | log(Present area) | 0.60 | 1.82 | 0.75 – 5.19 | 0.148 |
| Temporal | log(Present cost) | -0.92 | 0.40 | 0.10 – 1.31 | 0.093 |
| Structural | PC1_area | 0.31 | 1.36 | 0.61 – 1.90 | 0.244 |
| Structural | PC1_cost | -0.40 | 0.67 | 0.31 – 1.23 | 0.202 |

The Present model exhibited the same pattern with slightly reduced effect sizes (area: OR = 1.82; 95% CI: 0.75–5.19; cost: OR = 0.40; 95% CI: 0.10–1.31). Although these predictors did not reach conventional significance (p < 0.05), the consistency in effect direction and magnitude, combined with model-selection weights favoring the LGM (w = 0.53) over the Present (w = 0.50), indicates a clear historical legacy signal.

### H2 - Structural, time-integrated (PCA composites)

PCA effectively summarized cross-period gradients for area and connectivity. PC1_area captured a shared “large-across-time” signal (69% variance explained; loadings: LGM 0.74, Present 0.60, MH 0.35), while PC1_cost represented “low-isolation-across-time” (66% variance explained; loadings: LGM 0.71, Present 0.58, MH 0.41). The variance explained values correspond directly to the PC1 eigenvalues, and the reported loadings represent the eigenvector coefficients defining each axis.

The additive structural model received moderate support (AICc = 30.7; ΔAICc = 2.6; AUC = 0.725; CV-AUC = 0.533). As predicted, PC1_area had a positive effect on occupancy (β = 0.31; OR = 1.36; 95% CI: 0.81–2.30; p = 0.244), while PC1_cost had a negative effect (β = −0.40; OR = 0.67; 95% CI: 0.31–1.23; p = 0.202). These results suggest additive, compensatory effects of habitat extent and isolation across time, consistent with H2, though overall support was weaker than for the LGM-specific model.

### Temporal Trends in Habitat Configuration

Climatic suitability for *S. arizonensis* exhibited dynamic elevational and spatial shifts through time rather than a simple monotonic trend (Fig. 5). The suitability-weighted mean elevation increased from 1,643 m during the LGM to 1,792 m in the Mid-Holocene, declined to 1,723 m at Present, and is projected to rise again to 1,782 m under the Future scenario. Geographic centroids followed a similarly non-linear trajectory: from (−110.006°, 32.242°) during the LGM, shifting northwest to (−110.413°, 32.415°) in the MH, southwest to (−110.508°, 31.831°) at Present, and northwest again to (−110.613°, 32.028°) under Future climate.

**Figure 5.**
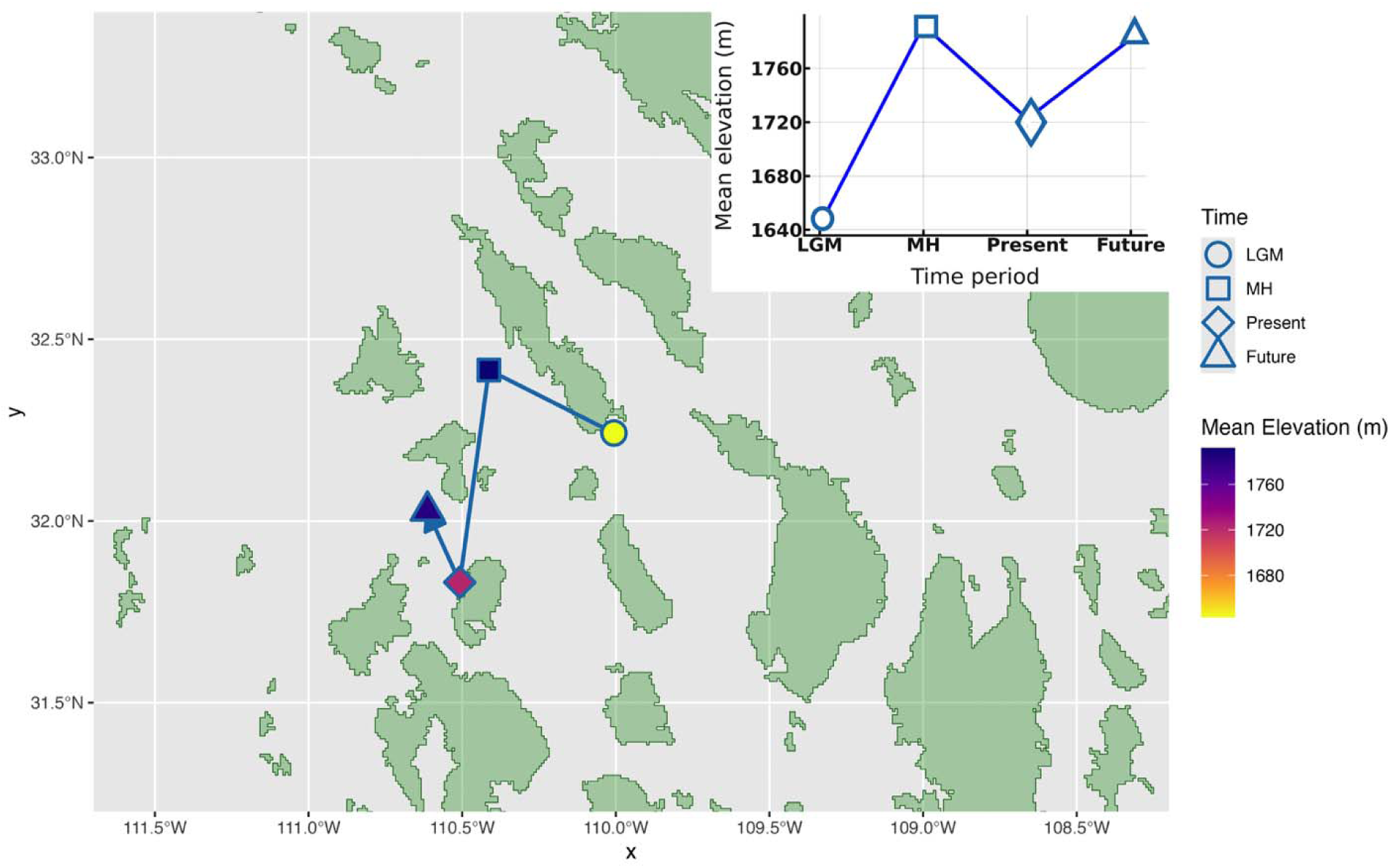
Geographic and elevational shifts in *Sciurus arizonensis* habitat centroids across four time periods. Symbols mark centroids (circle = LGM, triangle = Mid-Holocene, square = Present, diamond = 2070 projection) and are color-coded by mean elevation. Arrows trace the sequential movement of each centroid, with the inset line graph showing the corresponding change in suitability-weighted mean elevation. Centroids shift northwestward and upslope from the LGM to the Mid-Holocene, southwestward and downslope toward the Present, and northwestward and upslope again under the Future scenario. Sky-island terrain above 1,500 m is shaded green to provide geographic context.

## Discussion

### Pleistocene refugia and historical legacy

Our results suggest a legacy of long-term occupancy, in which the present distribution of *S. arizonensis* reflects habitat extent and connectivity during the LGM. Models incorporating LGM area and cost consistently outperformed those based on MH and slightly exceeded the predictive power of present-day models, underscoring an enduring influence of late-Pleistocene landscapes (Table 1). Independent paleoecological evidence, including packrat middens, pollen records, dynamic vegetation simulations, and regional genetic studies, suggests that valleys now separating the Madrean ranges supported continuous pine–oak and mixed conifer woodlands during the LGM (Van Devender et al. 1987; Betancourt et al. 1990; Spaulding 1990; Allen and Anderson 2000; Harrison and Prentice 2003; Brierley et al. 2020; Willyard et al. 2021). These low-elevation woodlands would have provided functional corridors for species such as *S. arizonensis.* Comparable patterns have been observed in other tree squirrels, where historical habitat stability predicts modern genetic diversity and population structure (Hernández, Fernández and Upham 2025), reinforcing the idea that long-term habitat configuration shapes both distribution and genetic differentiation across fragmented landscapes.

Although our models do not explicitly test extinction dynamics, the results suggest that populations established under glacial conditions have persisted to the present, consistent with long-term occupancy driven by historical connectivity rather than recent colonization. Moreover, the LGM and MH likely represent recurring phases within longer Pleistocene climatic cycles of habitat expansion and contraction (Byrne 2008; Meriño, Villalobos-Barrantes and Guerrero 2024). Thus, *S. arizonensis* populations may have been established well before the LGM, with successive climatic oscillations reinforcing the spatial legacy evident today.

Within the LGM model, larger refugial forest area increased occupancy probability, whereas higher historical travel cost decreased it. Although neither predictor reached conventional significance, their directions and magnitudes align with island-biogeographic expectations and broader evidence that Pleistocene refugia continue to shape modern biodiversity patterns (Hewitt 2000; McCormack et al. 2008; Love et al. 2023). Reduced woodland connectivity during the MH likely created a dispersal bottleneck, limiting recolonization after local extinctions and reinforcing historical constraints on present-day occupancy.

### Habitat area and current landscape effects

Present-day habitat area exhibited a positive, although weaker, association with *S. arizonensis* occupancy, while travel cost retained a negative effect. The Present model performed nearly as well as the LGM model (ΔAICc = 0.1) and achieved a slightly higher cross-validated AUC (0.733 vs. 0.717), suggesting that contemporary fragmentation still influences occupancy patterns. Together, these findings are consistent with a distributional disequilibrium scenario (Svenning and Skov 2004), in which some mountains remain unoccupied despite suitable current climate because they were historically disconnected. The PCA-based structural model, which condensed predictors across periods into orthogonal axes (PC1_area and PC1_cost), produced the expected directions of effect but explained less variance overall. This outcome reinforces that glacial refugial conditions were the dominant determinants of persistence, whereas modern habitat configuration now modulates occupancy at finer spatial scales.

### Climatic preferences and environmental filtering

Maxent response curves (Supplementary Data: SD4) indicate that *S. arizonensis* occupies a relatively narrow climatic niche. Climatic suitability peaks at moderate annual temperatures (BIO1) and intermediate precipitation levels of approximately 500–600 mm (BIO12), declining toward both hotter–drier and cooler–wetter extremes. Suitability also increases under moderate precipitation seasonality (BIO15) and higher driest-quarter rainfall (BIO17), reflecting sensitivity to both the amount and timing of moisture. These climatic thresholds are consistent with the species’ dependence on riparian corridors within pine–oak woodlands and its limited tolerance for arid conditions, where water availability and canopy cover likely buffer thermal and hydric stress (Frey et al. 2008; Cudworth and Koprowski 2011). Together, these results highlight a climatic filtering process that complements the historical-connectivity legacy identified above, jointly constraining the distribution of *S. arizonensis* across the Madrean Sky Islands.

### Constraint-Based Dynamic Island Biogeography (C-DIB)

Our findings align with the C-DIB framework (Burger et al. 2019), which emphasizes how historical connectivity and habitat area continue to shape present-day occupancy patterns. Within this framework, past isolation can lead to local extinctions and ecological hysteresis, where species distributions retain the imprint of former landscapes despite subsequent environmental change. In line with these predictions, historically isolated ranges in our study rarely regained *S. arizonensis* populations even where current climatic conditions appear suitable. This pattern underscores the enduring influence of historical constraints and highlights the C-DIB as a valuable extension of classical island biogeography theory for explaining persistence and absence in fragmented systems.

### Elevational and Geographic Reconfiguration of Climatically Suitable Habitat

Model projections reveal substantial restructuring of *S. arizonensis’* climatic niche since the LGM. The suitability-weighted mean elevation rose by roughly 150 m from the LGM (1,643 m) to the MH (1,792 m), declined to 1,723 m at Present, and is projected to rise again to about 1,782 m by the late century. These non-linear elevational changes parallel regional climatic trends, reflecting post-glacial warming followed by MH aridity (Van Devender et al. 1987; Freeman et al. 2018).

Spatially, the centroid of suitable habitat shifted northwest from the LGM to the Mid-Holocene, southwest from the Mid-Holocene to the Present, and modestly northward under future projections, reflecting temporal changes in regional moisture regimes (Metcalfe et al. 2015). The largest displacements occurred well after the initial colonization phase, indicating major habitat reconfiguration long after dispersal pathways had closed. Ecologically, this pattern suggests that many present populations now occupy only fragments of their ancestral elevational range, leaving them vulnerable to a “summit-trap” dynamic that heightens extinction risk (Haase et al. 2019). Conversely, projected downslope expansion of suitable climate under future scenarios points to potential habitat recovery, provided riparian corridors remain intact, emphasizing the importance of management strategies tailored to range-specific elevational responses.

### Biogeographic and Conservation Implications

The LGM model, integrating refugial area and dispersal cost, slightly outperformed the Present model, suggesting that populations established during glacial periods have persisted as habitats fragmented, whereas climatically suitable yet historically isolated ranges remain unoccupied.

This pattern fits a distributional disequilibrium framework in which past dispersal barriers continue to shape present-day distributions (Svenning and Skov 2004). This historical signal is consistent with the species’ life-history traits, limited dispersal, dependence on riparian corridors, and a slow reproductive rate (Hoffmeister 1986; Best and Riedel 1995), all of which constrain recolonization. Its persistence is further jeopardized by competition with the more generalist S*. aberti*, which can expand into *S. arizonensis* habitat following disturbance (Davis and Bissell 1989; Edelman and Koprowski 2005); recent evidence shows that the two species respond differently to post-fire landscapes, with *S. aberti* better exploiting burned areas (Ketcham et al. 2017). Additional biotic interactions may also contribute to the patchy occupancy observed across the Madrean region. For example, the restricted U.S. range of *S. nayaritensis* relative to its broader southern distribution may reflect historical competition or shifting range boundaries.

While our models primarily quantify area and connectivity, these ecological and behavioral interactions likely act as additional filters influencing persistence on otherwise suitable islands. Anthropogenic disturbances further compound these pressures. Degradation of riparian zones through logging, fire suppression, and grazing (Doumas et al. 2015) has reduced habitat quality and continuity. Increasingly severe wildfires have also caused widespread losses of montane forest cover; even when riparian corridors remain, the destruction of intervening upland forests can fragment metapopulations and isolate local populations.

Our analyses indicate that the current distribution of *S. arizonensis* across the Madrean Sky Islands still bears the imprint of Late Pleistocene connectivity: ranges that were either large or well-connected during the LGM remain occupied today, whereas historically isolated mountains are typically unoccupied even where suitable climate persists. This pattern reflects a state of distributional disequilibrium and aligns with predictions from the C-DIB framework, underscoring the importance of integrating palaeoecological legacies into conservation planning. Several conservation priorities emerge from these findings. Protecting large, contiguous habitat patches remains critical for long-term persistence, while restoring and maintaining riparian corridors will enhance functional connectivity among isolated populations, particularly for dispersal-limited species such as *S. arizonensis*. Because upland forests influence the continuity of these corridors, maintaining upland buffers is also essential to prevent metapopulation collapse. Moreover, each sky island should be managed as a discrete conservation unit, reflecting both its historical isolation and its potential for local adaptation (Hoffmeister 1986). Expanding native riparian habitats is also critical given ongoing climate-driven forest contraction (Steele and Koprowski 2003; Neary et al. 2005). Although assisted translocation may be appropriate for the most isolated ranges, such interventions should be guided by genomic assessments to avoid disrupting local adaptation, with natural connectivity enhancement remaining the preferred strategy (Aitken and Whitlock 2013; Chen et al. 2022). Continued ecological and genetic research will be essential to refine these approaches and strengthen resilience in *S. arizonensis* and other sky island taxa facing accelerating environmental change.

## Acknowledgement

We acknowledge the Burger and Ferguson labs at the University of Kentucky and the Constraint Based Dynamic Island Biogeography Working Group, including Rob Anderson, Trevor Fristoe, Kimberly Cook and Andrew Gaier for support and feedback throughout this study.

## Author Contributions

Binaya Adhikari: Conceptualization, Methodology, Data curation, Formal analysis, Investigation, Visualization, Validation, Writing – original draft, Writing – review & editing. Jesse M. Alston: Supervision, Validation, Writing – review & editing.

Joseph R. Burger: Conceptualization, Methodology, Supervision, Validation, Funding acquisition, Writing – review & editing.

## Supplementary Data

Supplementary Data SD1— Study area map focused to show occurrence records (GBIF) of four tree squirrel species across the Rocky Mountains and Colorado Plateau (mainland) and the Madrean Sky Islands. Points represent *Sciurus arizonensis* (blue), *S. nayaritensis* (red), *Tamiasciurus fremonti* (orange), and *S. aberti* (yellow); diamonds indicate preserved specimens and circles denote human observations. Labels mark introduced populations (*S. aberti*, “I”) and questionable localities (“?”). Background vegetation classes are derived from the LANDFIRE (LANDFIRE 2020) database and applied consistently across both U.S. and Mexican ranges: mixed conifer–oak (“conifer–hardwood,” dark green), conifer (light green), and riparian forest (blue). The inset shows the species’ broader ranges and study region. Photographs depict the four focal species.

Supplementary Data SD2— Bioclimatic predictors retained in the Maxnet climatic-suitability model for *Sciurus arizonensis*: Variance Inflation Factors (VIF), percentage contribution, and permutation importance were derived from the final Maxnet model (feature classes: LQHP; regularization multiplier β = 1). All predictors exhibited acceptable multicollinearity (VIF < 6). Relationships summarize the ecological response of *S. arizonensis* to each variable based on response curves, illustrating temperature and precipitation gradients that define the species’ climatic niche.

Supplementary Data SD3— Climatically suitable habitat patch maps for *Sciurus arizonensis* across four time periods: Last Glacial Maximum, Mid-Holocene, Present, and Future. Maps were generated using a 10th-percentile (p10) threshold applied to continuous Maxnet climatic-suitability outputs. White areas indicate suitable climate (above threshold), and black areas represent unsuitable conditions (below threshold). Red outlines delineate terrain above 1,500 m elevation, corresponding to the fixed sky island boundaries used for calculating total suitable area per island and period.

Supplementary Data SD4— Response curves from the selected Maxnet species distribution model (feature classes = LQPH; regularization multiplier β = 1) showing how key bioclimatic variables influence predicted climatic suitability for *Sciurus arizonensis*. The x-axes represent environmental values for each variable—BIO01 (Annual Mean Temperature), BIO07 (Temperature Annual Range), BIO12 (Annual Precipitation), BIO15 (Precipitation Seasonality), and BIO17 (Precipitation of the Driest Quarter)—while the y-axis shows the predicted probability of occurrence (cloglog output).

